# Late Miocene transformation of Mediterranean Sea biodiversity

**DOI:** 10.1101/2024.03.14.585031

**Authors:** Konstantina Agiadi, Niklas Hohmann, Elsa Gliozzi, Danae Thivaiou, Francesca R. Bosellini, Marco Taviani, Giovanni Bianucci, Alberto Collareta, Laurent Londeix, Costanza Faranda, Francesca Bulian, Efterpi Koskeridou, Francesca Lozar, Alan Maria Mancini, Stefano Dominici, Pierre Moissette, Ildefonso Bajo Campos, Enrico Borghi, George Iliopoulos, Assimina Antonarakou, George Kontakiotis, Evangelia Besiou, Stergios D. Zarkogiannis, Mathias Harzhauser, Francisco Javier Sierro, Marta Coll, Iuliana Vasiliev, Angelo Camerlenghi, Daniel García-Castellanos

## Abstract

Understanding deep-time marine biodiversity change under the combined effects of climate and connectivity changes is fundamental for predicting the impacts of modern climate change in semi-enclosed seas. We quantify the Late Miocene–Early Pliocene (11.63–3.6 Ma) taxonomic diversity of the Mediterranean Sea for calcareous nannoplankton, dinocysts, foraminifera, ostracods, corals, molluscs, bryozoans, echinoids, fishes, and marine mammals. During this time, marine biota was affected by global climate cooling and the restriction of the Mediterranean’s connection to the Atlantic Ocean that peaked with the Messinian Salinity Crisis. Although the net change in species richness from the Tortonian to the Zanclean varies by group, species turnover is greater than 30% in all cases. The results show clear perturbation already in the pre-evaporitic Messinian (7.25–5.97 Ma), with patterns differing among groups and sub-basins.

## Introduction

Climate and connectivity control the structure and functioning of marine ecosystems (*1*). Although this statement is supported by theory and observation, the response of the different groups of organisms to combined climate-connectivity changes remains unclear. The Mediterranean is a model marginal oceanic basin, whose ecosystem is profoundly altered by climate and connectivity changes today. A semi-enclosed basin, the Mediterranean is at present one of the places most impacted by climate warming (*2*), as well as by impressive rates of invasion by alien species from the Indo-Pacific realm after the opening of the Suez Canal in 1869 (*3*). Under such dynamic conditions, it is challenging to predict community and ecosystem future states, and it is therefore important to look to the geological past, for periods of extreme environmental change. In this respect, the Late Miocene Mediterranean Sea is the ideal setting.

The Late Miocene (11.63–5.33 Ma) was a pivotal time for the Mediterranean marine biota. The global climate cooling (*4*) and the basin’s disconnection from the Atlantic Ocean during the Messinian Salinity Crisis (MSC; *5*, *6*) led to extreme environmental conditions in the basin. However, the impact of this ecological crisis on marine biodiversity has never been systematically studied. Under the climate cooling trend, a series of precursor events led to the MSC, the first being the stepwise restriction of the basin’s connections to the world ocean, which started already in the Earliest Messinian (*7*, *8*). High-stress conditions for marine organisms have already been reported immediately after the Earliest Messinian (7.17 Ma; *9*, *10*). Gradually, the restriction of the Rifian and Betic corridors (in present-day North Morocco and South Spain, respectively) led to increased salinity, water-column stratification, and episodes of dysoxia on the sea bottom (*11*, *12*). The first MSC evaporites were deposited on the Mediterranean marginal basins at 5.97 Ma, although Zachariasse and Lourens (*13*) suggested that the MSC started already at 6 Ma, at least in the eastern Mediterranean (eMed). There is general consensus that a one-way periodic connection between the Mediterranean and the Atlantic was maintained at least during the initial stage of the MSC (5.97–5.66 Ma; *8*, *14*, *15*). Even more extreme conditions occurred during the second MSC stage (5.6–5.55 Ma; *16*), as evidenced by the kilometer-thick deposits of salt found throughout and even in the deep parts of the Mediterranean. In the final stage of the MSC, periodic alternations of gypsum and marls (5.55–5.42 Ma) followed by the brackish ‘Lago Mare’ deposits (5.42–5.33 Ma) reflect increased freshwater influx to the basin possibly from the Paratethys in the North (*17*), although the Paratethyan-inflow hypothesis has been contested (*18*). Normal marine conditions were established once more in the Mediterranean at the base of the Zanclean at 5.33 Ma (*19*), after the restoration of the connection with the Atlantic Ocean (*20*).

The literature is full of hypotheses on the magnitude of the MSC repercussions on the different groups of organisms (e.g., *21*), but these are mostly based on incomplete and still uncertain scenarios about the MSC evolution, as well as on the intuitive assumption that such a paleoenvironmental perturbation must have caused a ‘major’ change in the marine biota. Having prevailed for many decades now, this assumption has leaked from paleontology and geosciences to biological sciences, with numerous papers referring to it as a fact (e.g., *22*), instead of what it truly is, only an assumption. However, some studies have opposed this view: for example, Néraudeau et al. (*23*) stated that the “Messinian desiccation was not a drastic event for irregular echinoids”, and Goubert et al. (*24*), studying the benthic foraminifera assemblages at Los Yesos (Sorbas Basin, Spain), supported that “the MSC is not associated with a biological crisis”. Moreover, Monegatti and Raffi (*25*) reported an important impact of the MSC on marine gastropods, but a very small impact on bivalves, highlighting the need for a more in-depth, integrated ecosystem-based assessment.

In this study, we analyze a recently revised Tortonian–Zanclean marine fossil record of calcareous nannoplankton, dinocysts, planktic and benthic foraminifera, ostracods, corals, bivalves, gastropods, bryozoans, echinoids, bony fishes, sharks and marine mammals, from the western Mediterranean (wMed), the eastern Mediterranean (eMed) and the Po Plain– Northern Adriatic (PoA) region (*26*; Figs. 1 and 2) to obtain evidence of changes in the taxonomic diversity of the Mediterranean marine biota that took place from the time of the initiation of the Mediterranean–Atlantic gateway restriction in the Late Tortonian (*27*), until the establishment of a fully marine ecosystem in the Zanclean. Investigations of the fossil record from the MSC beds has been presented elsewhere, indicating that stenohaline marine organisms appeared in various levels (*28*, *29*). However, in our investigation, we exclude the interval corresponding to the MSC because the fossil record recovered from the MSC beds is limited compared to the record before and after the crisis, and it cannot be used therefore for the present biodiversity analysis. Instead, here we take a step back and evaluate the biodiversity change by comparing the Tortonian to the pre-evaporitic Messinian and to the Zanclean fossil record of these groups. Taxonomic diversity is examined by calculating four diversity metrics: richness, total dissimilarity (Sørensen index), dissimilarity due to turnover (Simpson index), and nestedness (*30*). Our results demonstrate the different impact that changes in basin connectivity and climate had on the composition of the marine assemblages of the different groups, and highlight research questions that remain open.

**Figure 1.**
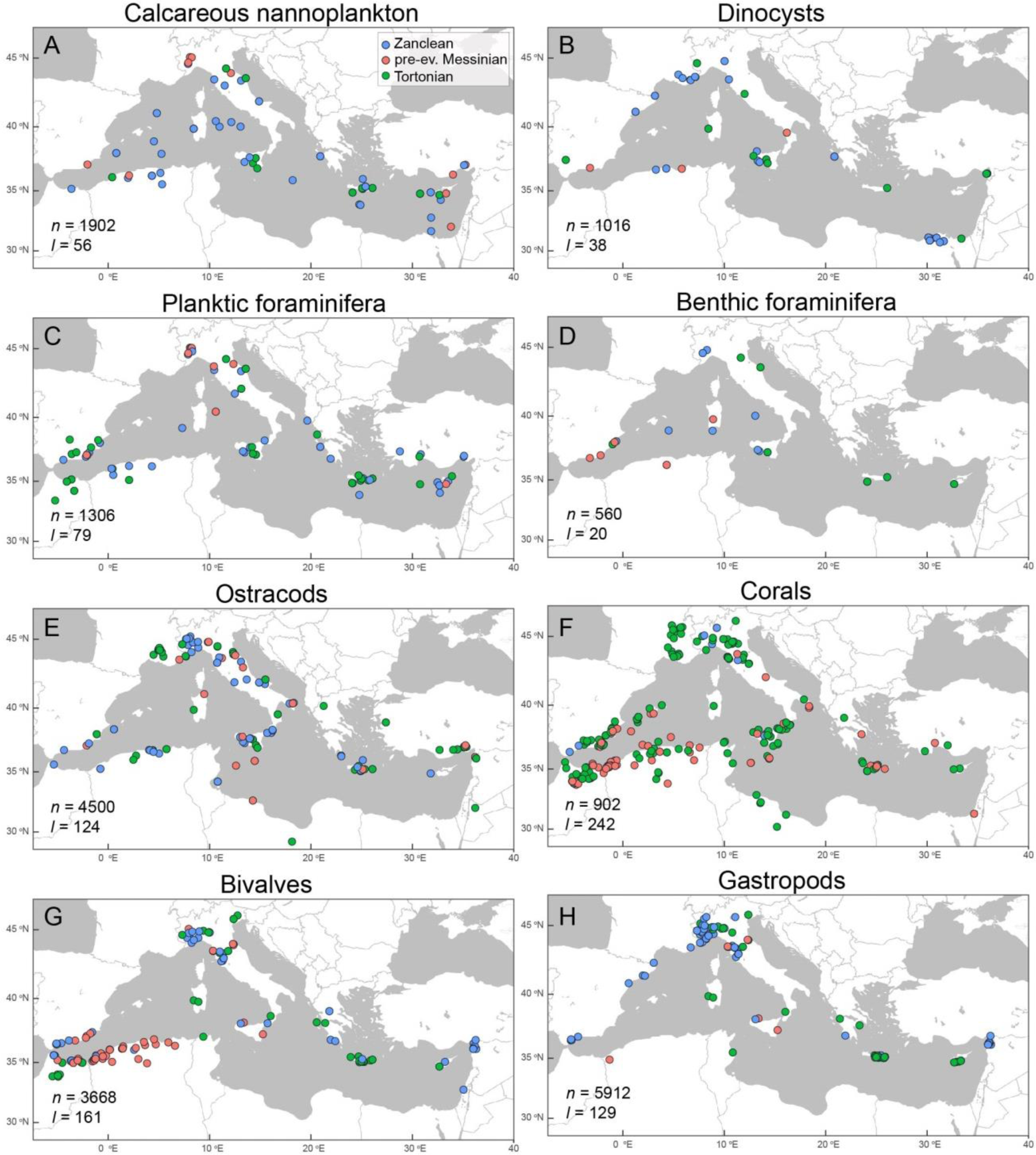
Mediterranean map with the localities included in this record. (*26*). A. Calcareous nannoplankton, B. Dinocysts, C. Planktic foraminifera, D. Benthic foraminifera, E. Ostracods, F. Corals, G. Bivalves, H. Gastropods. The maps were produced using ggmap (*76*). *n* indicates the number of occurrences in the dataset, and *l* indicated the number of localities.

**Figure 2.**
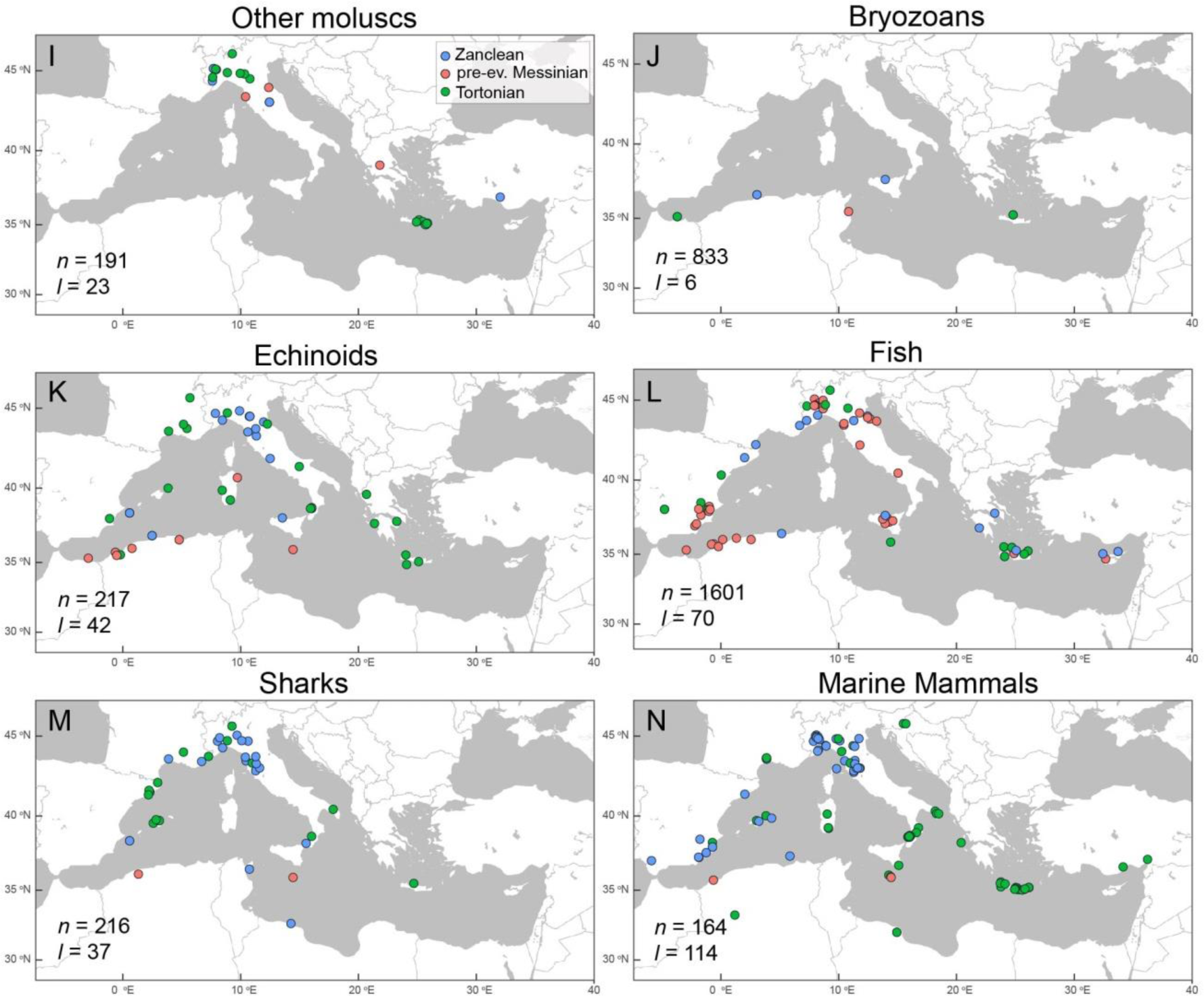
Mediterranean map with the localities included in this record. (*26*). I. Other molluscs (scaphopods, chitons, cephalopods), J. Bryozoans, K. Echinoids, L. Fish, M. Sharks, N. Marine mammals.

## Results

The Late Miocene–Early Pliocene Mediterranean species richness patterns vary greatly between taxonomic groups (Fig. 3) and sub-basins (supplement). Species richness decreased from the Tortonian to the pre-evaporitic Messinian for calcareous nannoplankton (by 11.6%; driven by wMed, Fig. S1), dinocysts (5.3%, also driven by wMed, Fig. S2), planktic foraminifera (23.4%; mostly in wMed, Fig. S3), corals (60%; although N=18 through), bivalves (35.3%; large decrease in wMed, but increase in eMed, Fig. S5), echinoids (10.3%; driven by wMed where most data come from, Fig. 2), and bony fishes (21.8%; driven by PoA, Fig. S7). Ostracod species richness remained at the same levels from the Tortonian to the Messinian, but this is driven by the PoA record, while richness increased in both wMed and eMed (Fig. S4). In contrast, species richness increased from the Tortonian to the Messinian for benthic foraminifera (18.4%), gastropods (6.3%; driven by PoA), and bryozoans (16.9%). From the Messinian to the Zanclean, species richness remained approximately the same for calcareous nannoplankton (increases in wMed, but decreases in eMed, Fig. S1), planktic and benthic foraminifera, gastropods, and bryozoans, whereas there is a decrease in richness for dinocysts (19.1%; driven by eMed, Fig. S2), ostracods (45.9%; also mostly driven by eMed, Fig. S4), and echinoids (7.4%). Species richness increased from the Messinian to the Zanclean for corals (41.2%; only cold-water corals, mostly azooxanthellate in the Zanclean, *33*), bivalves (32.1%), and bony fishes (19.8%).

**Figure 3.**
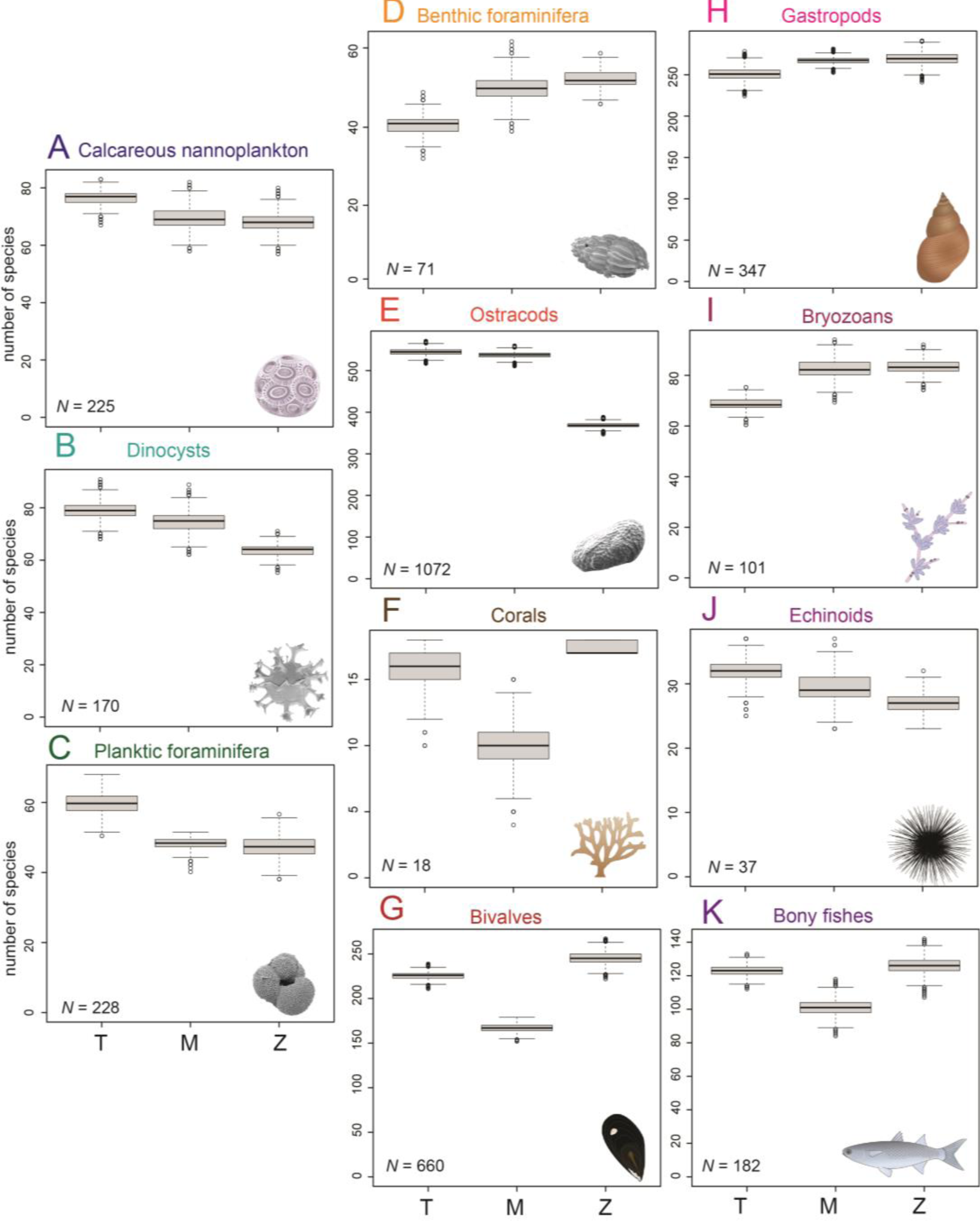
Changes in species richness of the Mediterranean Sea biota from the Late Miocene to the Early Pliocene by group of organisms. X-axes shows intervals: Tortonian (T), pre-evaporitic Messinian (M), and Zanclean (Z). Symbols for organisms obtained from the Integration and Application Network (ian.umces.edu/media-library). *N* indicates the number of occurrences after subsampling to 80% of the smallest sample, ten thousand times. The bold line indicates the median value, the box corresponds to the quartiles, whiskers are quartiles plus/minus 1.5 times the interquartile range.

Species turnover always contributes more than nestedness to the total dissimilarity, in all comparisons (Fig. 4). Total dissimilarity and species turnover are higher in the Messinian- versus-Zanclean comparisons of all groups, whereas nestedness is highest in the Tortonian- versus-pre-evaporitic Messinian comparisons. Total dissimilarity between the Tortonian and pre-evaporitic Messinian records exceeds 50% for corals (76.0%), gastropods (72.8%; driven by PoA), bryozoans (53.9%), echinoids (77.4%), bony fishes (70.6%), whereas it is less than 50% for calcareous nannoplankton (25.2%), dinocysts (35.0%), planktic (30.0%) and benthic foraminifera (42.2%), ostracods (37.8%), and bivalves (43.0%). In the Messinian-versus- Zanclean comparisons, total dissimilarity is greater than 50% in benthic foraminifera (61.6%), ostracods (68.7%), corals (100%), gastropods (77.2%), bryozoans (64.9%), echinoids (67.3%), and bony fishes (83.7%). Only calcareous nannoplankton (31.4%), dinocysts (45.5%), planktic foraminifera (40.7%), and bivalves (48.4%), maintain lower than 50% dissimilarities.

**Figure 4.**
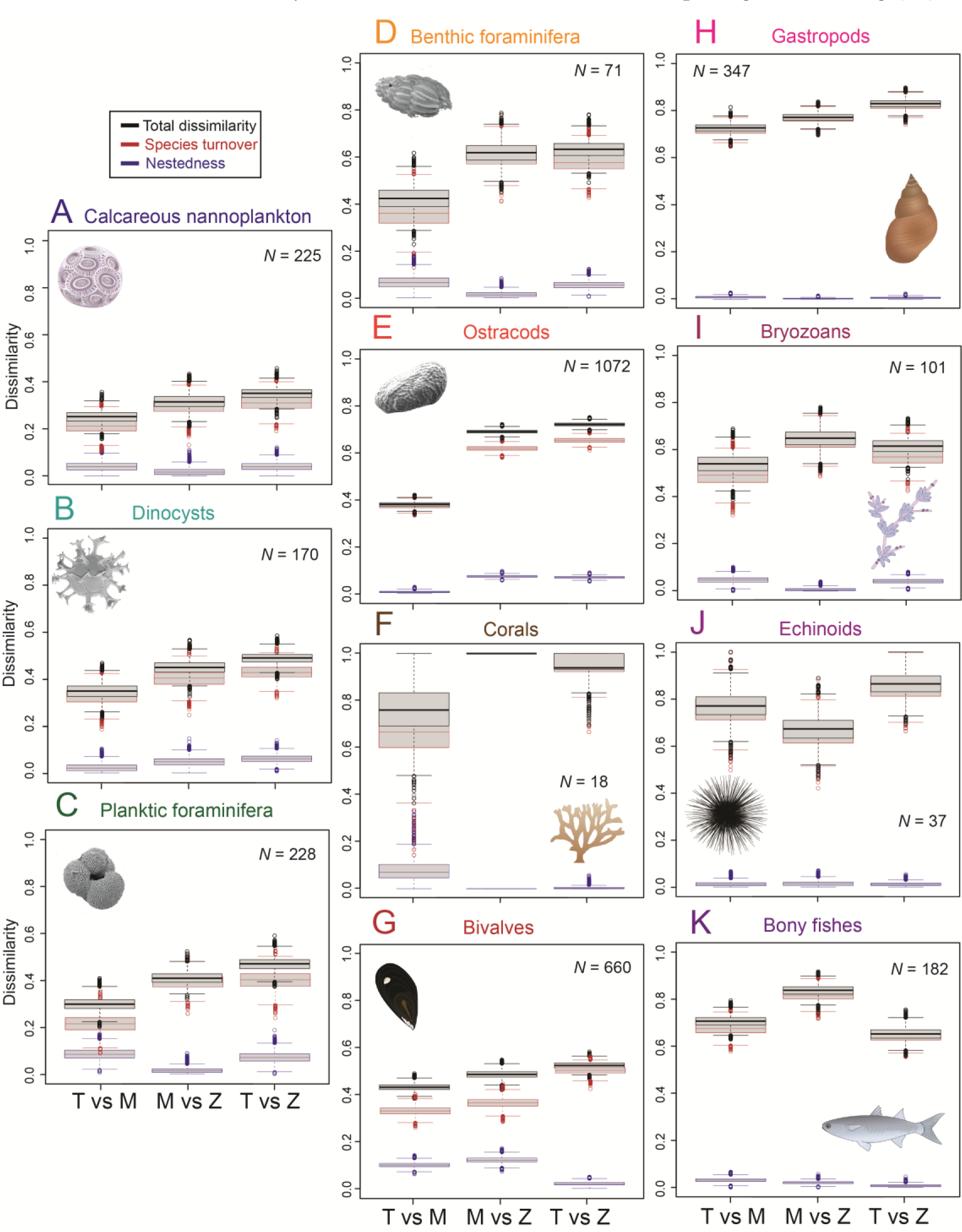
Temporal change: dissimilarities between the Tortonian (T), pre-evaporitic Messinian (M) and Zanclean (Z) biodiversities of the Mediterranean for each group of organisms. Black: total dissimilarity; Red: species turnover; Blue: nestedness.

Finally, the net effect (Tortonian-versus-Zanclean comparisons) varies among groups, with species turnover of 31.0% for calcareous nannoplankton, 41.7% for dinocysts, 40.0% for planktic and 57.5% for benthic foraminifera, 65.0% for ostracods, and 93.8% corals. These values of species turnover are at the levels achieved during the pre-evaporitic Messinian. Gastropods and echinoids exhibit even higher species turnover in the Tortonian-versus- Zanclean (82.5% and 85.2%) than in the Messinian-versus Zanclean comparison (77.0% and 65.4%, respectively). The Tortonian-versus-Zanclean faunas of bryozoans (61.5%) and bony fishes (65.2%) are more similar than the Messinian-versus-Zanclean faunas (64.7% and 83.7%, respectively).

## Discussion

### Climatic-related triggers and connectivity

The Late Miocene–Early Pliocene Mediterranean marine biota resulted from the interplay between global climatic cooling and changes in marine connectivity within and beyond the Mediterranean Sea. The cooling directly affected temperature-sensitive organisms such as the tropical reef-building z-corals and their associated faunas (reef fishes and sharks) and bryozoans leading to local extinction of vast populations, particularly in the eMed (*31*).

Additionally, the decrease of water temperatures in the Mediterranean allowed boreal species to expand their distribution to the basin during the Messinian, while strongly thermophilic Tethyan relic species disappeared. Monegatti and Raffi (*25*) noted that the MSC caused regional mass disappearances of molluscs, but only a limited number of actual extinctions, and that the greatest Messinian extinctions took place in the Atlantic Ocean and were triggered by the TG22, TG20, TG14, and TG12 glacials during the MSC. In the Zanclean, the establishment of psychrospheric water masses in the Atlantic further exacerbated this impact. For example, the great white (*Carcharodon carcharias*) and the blue shark (*Prionace glauca*) first appeared globally at the Miocene/Pliocene boundary (*32*) and in the Mediterranean after the MSC (*33*). Shallow-water tropical coral reefs dominated by colonial zooxanthellate scleractinian corals (z-corals) exhibited low diversity in the Late Miocene and almost disappeared from the Mediterranean after the MSC (*34*), accounting for the observed drop in species richness from the Tortonian to the Messinian (Fig. 3F) and the high species turnover between the pre- and post-MSC coral faunas (Fig. 4F). However, the Mediterranean coral species richness fully recovered in the Zanclean (Fig. 3F) due to the introduction of azooxanthellate (az-) coral species (mostly solitary deep-water corals). The MSC played a crucial role in the local extinction of shallow-water coral (z-coral) reefs, but it was not the main driver (*31*, *35*). Z-corals, as tropical reef corals, are highly sensitive to temperature. The marked decrease in their diversity actually started well before the MSC, in the late Middle Miocene, with the gradual northwards shift of the Mediterranean region outside the tropical belt, the closure of the seaway with the Indo-Pacific and the subsequent global cooling (*35*).

Whether climate or the water-column stratification and increased bottom-water salinity in the Messinian drove populations to seek refuge in the Eastern Atlantic off the western coast of Africa is unclear. The Pliocene survival of Mediterranean endemics taking refuge in the Atlantic during the MSC, and then repopulating the Mediterranean after the crisis, has been postulated by many researchers (*36*). Indeed, considering that the overall species turnover between Tortonian and Zanclean ostracod faunas is greater than that between Tortonian and Messinian (Fig. 4E), it seems likely that the Zanclean saw a return of some ostracod species that were present in the Mediterranean during the Tortonian but not the Messinian. The Atlantic part of the Rif, as well as southern Portugal and SW Spain may have been such Messinian refugees (*25*). Echinoids taking refuge in the North Atlantic and then repopulating the basin at the beginning of the Pliocene has been suggested (e.g., *23*), but is not fully supported by the fossil records of the Eastern Atlantic (e.g., *37*) show similar taxonomic composition but lower diversity than their coeval communities in the Mediterranean basin. This observation would support the hypothesis of Booth-Rea et al. (*38*) that the West Alboran Basin was an open-marine refuge for Mediterranean taxa that repopulated the sea after the MSC, connected to the deep restricted Mediterranean basin through a sill at the Alboran volcanic arc archipelago. However, Bulian et al. (*10*) demonstrated that the deep sea community of the West Alboran Basin (along with the rest of the Mediterranean) was disconnected from that of the Atlantic already from 7.17 Ma. Therefore, this area could not have functioned as a marine refuge for Mediterranean endemics during the MSC. Moreover, migrations to Eastern Atlantic refuges, although possible for a few species of dinocysts, ostracods, bryozoans, bivalves, gastropods, corals and fishes, are not supported by the nestedness between the pre- and post-MSC faunas which is consistently below 10% (Fig. 4). This percentage is better explained by the re-establishment of species that were already present in the Atlantic before the crisis.

The MSC precursor events created intervals of salinity fluctuations, bottom-water deoxygenation and stratification of the water column (*12*, *39*), conditions that gradually became extreme for most species inhabiting the basin during the Messinian. Most planktonic groups show a clear response, their diversity dropping in the pre-evaporitic Messinian. For example, calcareous nannoplankton and planktic foraminifera decreased in species richness in the wMed during the Tortonian/Messinian transition (Figs. S1 & S2), and this has been attributed to eutrophication as a result of the gateways’ restriction (*10*, *40*). In contrast, calcareous nannoplankton species richness remains quite unchanged through time in the eMed (Fig. S1), probably because the eMed was more affected by nutrient input provided through continental runoff (*41*), leading to a slightly more diverse assemblage already in the Tortonian, and so the effect of the restriction was not as strong in the eMed. On the other hand, the intra-Messinian salinity fluctuations strongly impacted plankton biodiversity in the eMed (*39*). Regarding benthos, eutrophication and bottom-water deoxygenation negatively affected benthic foraminifera species richness during the Early Messinian (*9*, *10*), but the change depicted in the deep bottom environments is not captured in the overall record (Fig. 3), because of the appearance of shallow-water species (*42*). Similarly, littoral ostracod faunas were not at all impoverished, but bathyal ostracod assemblages were affected by the gateway restriction and the changes in water circulation (*43*). Deep-water coral communities (both colonial and solitary) and az-corals maintained their diversity in the Late Miocene and further increased it in the Pliocene and Pleistocene (*31*). A decoupling of the responses of shallow and deep-water corals may have occurred within the pre-evaporitic Messinian, and the revised coral dataset (*26*) confirms this hypothesis, highlighting a drastic modification of the scleractinian coral pool of the Mediterranean during the Late Miocene and Early Pliocene due to the loss of the reef-building and shallow-water z-coral assemblages.

Ecophenotypic adaptations were an additional impact of the pre-evaporitic Messinian environmental stress. Although the diversity of Sirenia did not change, they present a characteristic example of morphological changes due to the early Messinian restriction and change in the Mediterranean marine ecosystem (*44*). Sirenians ate seagrasses, and their tusk enamel δ^13^C before and after MSC reflected a shift in seagrass associations from *Posidonia oceanica* meadows in the Tortonian to more euryhaline forms with larger rhizomes like *Zostera* in the Messinian. Large-tusked sirenian species ate seagrass species with large rhizomes, which they extracted from the sediment, whereas species with small tusks ate leaves and shallow rhizomes (*45*). The dwarf size and large tusks of the endemic Mediterranean species *Metaxytherium serresii* are considered ecophenotypic adaptations due to the restriction of the habitat space and the change in the available food resources in the Messinian, which were reversed after the MSC, leading to the establishment of *M. subappenicum* in the Pliocene (*44*).

The high turnover and the low nestedness in all groups before and after the MSC (Fig. 4) reflect the importance of endemism in forming the Mediterranean biota. Oceanic endemic species were driven by the restriction of the connection with the Atlantic oceanic pool, whereas the Messinian neritic faunas were strongly affected by the establishment of the Paratethys–Mediterranean connection at 6.1 Ma that enabled the migration of species from the Paratethys to the Mediterranean and the evolution of new endemic species (*46*). Overall, the changing conditions throughout the Messinian triggered the appearance of new endemic species in the case for benthic foraminifera, gastropods, and bryozoans, whose species richness increases from the Tortonian to the Messinian (Fig. 3). Even so, species turnover is greatest for all groups between the Messinian and the Zanclean. Only for the planktonic groups, dissimilarity is below 50% between all comparisons (Fig. 4) because the Mediterranean and Atlantic surface waters were well-connected, and endemism is generally low in these groups. Nevertheless, both pre- to post-MSC comparisons demonstrate higher species turnover than the comparison between the Tortonian and Messinian plankton communities. For bivalves and gastropods, habitat fragmentation in the Zanclean due to the separation of the Adriatic from the Tyrrhenian Seas contributed to new endemics, thus increasing turnover (Fig. 4H and 4I). In contrast, for ostracods, the re-colonization from the Atlantic at the end of the MSC took place gradually, with only a few bathyal, opportunistic species recorded in the basal Zanclean (*47*). Possibly, new littoral endemic Mediterranean ostracod species also appeared later, in the Piacenzian, which would explain the low Zanclean species richness in comparison to the Tortonian and pre-evaporitic Messinian ones (Fig. 3E). Moreover, the Zanclean gastropod record is enriched with small-sized forms (e.g, fissurellids, trochids, rissoids and cerithiids), whose absence in the Late Miocene may be attributed to preferential loss by dissolution, difference in study design (few sieved samples are available) and difficulty in extraction of the specimens due to increased cementation.

Some bioevents may be attributed to first-order regional-scale oceanographic episodes. Overall, biotic immigration and isolation cycles have been shown to increase global biodiversity (*48*). This is not supported by a scenario where global extinctions of mollusc species were facilitated by the MSC that prevented these organisms from seeking refuge in the Mediterranean during Messinian glacials (e.g., *25*). The most striking signal comes from the presence in the Early Pliocene of taxa indicative of deep psychrospheric water masses from the Northern Adriatic, reflecting an estuarine circulation regime at Gibraltar. Specifically, the shark remains in the lower Zanclean sediments belong to species, which on the whole comprise a deep-water paleocommunity depicting a high degree of “oceanization” of Mediterranean Sea. At the same time, the high diversity and the turnover from archaic (i.e. Eurhinodelphinidae and Squalodontidae) to modern (i.e. Balaeonopteridae, Kogiidae and Pontoporiidae) families of cetaceans coincides with the Middle–Late Miocene global diversity peak that may be related to high, diatom-driven marine productivity combined with climatic change (*49*). Printed over a gradual declining trend associated with climate cooling, the observed negative peak of the global marine mammal diversity in the Messinian could have been caused by the MSC (*50*) or it could be an artifact due to lack data for this age for the Mediterranean Basin (*51*). Several cases support the dominance of global rather than regional patterns determining Mediterranean marine mammal diversity. For example, oceanic dolphins (Delphinidae) first appeared at the beginning of the Pliocene in the Mediterranean (*49*), but also in the North Atlantic. Nevertheless, within the Mediterranean, oceanic dolphins were more diverse, which could be due to the increased research effort in this region.

Alternatively, this high diversity combined with an apparently high degree of species- and genus-level endemicity could be interpreted as an indirect consequence of the MSC: delphinids may have recolonized the Mediterranean at the beginning of the Pliocene from the Atlantic Ocean, occupying the ecological niches available after the extinction/extirpation of the Miocene cetacean fauna (*49*).

## Limitations

The distribution of occurrences (Figs. 1 and 2) is highly uneven both geographically and in terms of sedimentary facies representation, leading to sampling bias (*52*, *53*). First, the fossil record analyzed here (*26*) was derived from paleontological studies conducted since the 19^th^ and beginning of the 20^th^ century, but these were constrained by the socio-economic and political conditions, favoring sampling in the European Mediterranean countries. Secondly, biodiversity estimated from the paleontological record depends on the volume and area of the sedimentary rocks exposed and potentially accessible for studies of their fossil content (*54*). This is in turn additionally related to the extent and method of sampling, which can also lead to large biases. Notably, in addition to the inherent sampling bias in paleobiodiversity studies (*55*), sampling approaches and methods changed drastically during the last 120 years. For large animals for example, such as marine mammals and fishes, selective sampling in the early 20^th^ century focused only on large specimens, easy to retrieve and handle, and often disregarding smaller fossils (*56*). Additionally, environmental factors (such as currents and turbidity, pH, hydrostatic pressure, sedimentation rate etc.) largely determine the quality (and quantity) of the fossil record through taphonomic preservation (*57*). For instance, an important part of the recorded Zanclean diversity of bivalves and gastropods is contributed by small-sized aragonitic forms, which could be underrepresented in the Late Miocene collections due to preservation and/or sampling biases, due to the difference in facies representation before and after the MSC in the Mediterranean. Marl and sand facies, which facilitate the preservation of small taxa, are common in the Zanclean (e.g., *58*) but nearly absent in the Late Miocene. Additionally, coral reef environments, which formed major parts of the Late Miocene Mediterranean coasts but became absent in the Zanclean, exhibited high cementation and aragonic shells were easily dissolved there (e.g., *52*). Finally, certain records are simply inaccessible to scientists for practical reasons: for example, the sediment cores obtained by deep-sea drilling expeditions in the Mediterranean yielded important records of microfossil occurrences for the Pliocene, but they could neither be used to obtain information about larger animals because of their dimensions, nor could they provide pre-evaporitic records. Consequently, for individual groups, the occurrences are often unevenly distributed between the three time-intervals, leaving intervals with very few or even no occurrences.

For some groups, such as the ostracods, the available data cover all intervals, regions, and environmental settings adequately. However, there are several examples where a combination of the above biases has led to unevenly distributed fossil records. Even in the few areas with continuous composite stratigraphic sequences, such as Sicily and the Nile Delta, the diversity patterns through time within individual groups can be very different. For example, in the case of dinoflagellate cysts, sampled with the same methodology throughout the Tortonian– Zanclean stratigraphic interval, species richness is highest during the Messinian in the record of the Nile Delta, but lowest in Sicily record, compared to the Tortonian and Zanclean. The rarefaction method employed in the calculation of diversity indices to (at least partly) overcome these sampling biases does not allow us to quantitatively evaluate several records due to this lack of sufficient number of occurrences (arbitrarily set at 15 occurrences in the present analysis) in one or more of the time intervals for a given taxonomic group. Groups with few occurrences within an interval (Figs. 1 and 2; *26*) should be targeted for further systematic paleontological studies.

### Ecologic, oceanographic and climatic implications of biodiversity change

The biodiversity changes, which are quantified here at the taxonomic level and based on the current status of the fossil record, point towards an ecological crisis in the Mediterranean during the Messinian, even before and peaking at the MSC. Although controversy is still high on the true nature and evolution of the MSC events (e.g. *59*, *60*), direct and indirect ecosystem consequences of the MSC cannot be denied.

To advance beyond these outcomes, future research should explore the implications of these changes for food webs, ecosystem structure and function, and for biogeochemical cycles. Functional diversity refers to the traits and niches filled by species, which essentially control how diversity influences the functioning of the ecosystem (*61*). For example, communities with the same species richness (a measure of taxonomic diversity) may include species with vastly different traits (e.g., pelagic versus demersal lifestyle etc.), who occupy a different ecological niche in the ecosystem, thus forming very different ecosystems. It is possible that the MSC resulted in or facilitated changes in functional, in addition to taxonomic, diversity in the Mediterranean marine ecosystem. The most characteristic is the case of corals, where tropical, reef-building (zooxanthellate) corals disappeared completely from the Mediterranean after the MSC (*31*). Furthermore, the functional composition of the marine biota determines the structure of the food web and thus the flow of energy and nutrients. Critical intervals, such as the Messinian for the Mediterranean Sea, involve perturbations of the biogeochemical cycles. Another possibility is that the Messinian decrease in mesopelagic fish diversity may have led to a decline in carbon export efficiency. These and other hypotheses should be tested, which would have important implications for the subsequent evolution of the Mediterranean environment.

## Materials and Methods

### Experimental design

The dataset that was used contains 22988 fossil occurrences of 4959 species, including some occurrences which have been considered reworked (*26*). We excluded these from the present analysis.

In order to examine the biodiversity changes observed for each group of organisms through time within Mediterranean regions, we distinguished the fossil localities into three regions based on their paleogeographic placement in the Western Mediterranean (wMed), the Eastern Mediterranean (eMed), and the Po Plain–Northern Adriatic (PoA), following the currently accepted paleogeographic data (*62*, *63*). The PoA region developed as a paleoceanographic sub-basin of the Mediterranean in the Tortonian–Early Messinian, as evidenced by its distinct strontium isotopic signature (*64*) and the absence of halite deposits. We placed the Tortonian records from Piedmont and the Po Plain within the PoA region as well, attempting to test the hypothesis that this region held an already distinct marine biota in the Tortonian. Based on the Late Miocene paleogeographic evolution of Calabria and Sicily (*65–67*), we included the records from these areas as part of the eMed, since the marine connection with the wMed was located near its present location, possibly along present southern Sicily (*68*) or at the Sicily Channel (Malta Plateau; *69*).

### Statistical analysis

Biodiversity can be evaluated across scales by evaluating alpha, beta, and gamma diversity (*70*). Alpha diversity refers to species richness, which is the number of species within a community. Beta diversity represents the amount of differentiation between communities, which may be due to: a) species turnover, which is the replacement of species by novel (and different) ones, independent of species richness; and b) nestedness, resulting from species loss through extinction (*71*). Nestedness, as an independent property of communities within an ecosystem, shows the difference between the faunas compared in terms of loss of species: when two faunas exhibit low nestedness (i.e., they are not nested) one is not a subset of the other (*72*). Gamma diversity refers to the total biodiversity across a larger geographic area (e.g., *73*).

Here, we calculated at species level: 1) alpha diversity (richness = number of species or genera) in the Tortonian, the pre-evaporitic Messinian, and the Zanclean; and 2) beta diversity comparing the different time intervals for the entire Mediterranean and the three regions (*74*). We calculated the metrics: richness, Søerensen index (total dissimilarity between the compared intervals), Simpson index (dissimilarity due to species turnover), and nestedness (*30*, *72*). To overcome potential bias in our biodiversity estimates due to the differences in sampling effort between the regions and intervals, we employed an 80% rarefaction, which has been shown to produce robust results (*75*). In practice, we subsampled the record of the intervals that are compared to the 80% of the lowest number of occurrences between them, and we used the subsamples for calculating the diversity indices. Subsampling was repeated 10000 times.

The data for sharks and marine mammals were not analyzed statistically separately given the limited number of recorded specimens and taxa and because cetaceans were mostly represented by poorly preserved diagnostic material that did not allow determination below the family level.

## Acknowledgments

This research was funded in whole, or in part, by the Austrian Science Fund (FWF) Grant DOI 10.55776/V986. For the purpose of open access, the author has applied a CC BY public copyright license to any Author Accepted Manuscript version arising from this submission. The authors would like to thank Jan Steger for reviewing an early version of this paper. This is Ismar-CNR, Bologna, scientific contribution n. 2091.

## Funding

Austrian Science Fund (FWF) grant V986-N (KA)

COST Action CA15103 (KA, DT, FBu, FL, AMM, GK, EB, SDZ, MH, FJSS, IV, ACa, DGC)

MIUR-Italy, Department of Excellence grant, Article 1, Paragraph 337, law 232/2016 (EG, CF)

‘Severo Ochoa Centre of Excellence’ accreditation (CEX2019-000928-S) (MC)

European Commission through ITN SaltGiant (Horizon2020-765256) (FBu, FJS, AC, DGC)

## Author contributions

Conceptualization: KA, ACa, DGC Data curation: KA, NH

Funding acquisition: KA, MC, ACa, DGC

Investigation: KA, NH, EG, DT, FBu, MT, GB, ACo, LL, CF, FBo, EK, FL, AMM, SD, PM, IBC, EB, GI, AA, GK, EB, SDZ, MH, FJSS

Formal analysis: NH Methodology: KA, NH Project administration: KA

Writing – original draft preparation: KA

Writing – review & editing: NH, EG, DT, FRB, MT, GB, ACo, LL, CF, FB, EK, FL, AMM, SD, PM, IBC, EB, GI, AA, GK, EB, SDZ, MH, FJSS, MC, IV, ACa, DGC

## Data and materials availability

All data are available in the main text or the supplementary materials. The dataset used for the present analysis is publically available under a CC BY license at https://zenodo.org/records/10782429. All code used in the analysis is available at https://zenodo.org/doi/10.5281/zenodo.10787996.

## Supplementary materials

### Additional background and results by taxonomic group

#### Dinocysts

The protoperidinioid/ gonyaulacoid cysts ratio in the record (Table S1) suggests higher productivity and hydrological surface circulation in the wMed during the Tortonian, than in the pre-evaporitic Messinian, but lower during the Zanclean, probably reflecting the restriction of the Tortonian wide connections with the Atlantic. The ratio of protoperidinioid over gonyaulacoid cysts is an indicator of the abundance of cysts produced by heterotrophic versus autotrophic dinoflagellates (*77*), and it is considered a measure of primary productivity. However, information on preservation can also be contained in this index (*78*), as well as bias due to the sample processing method (*79*). Moreover, the cysts of protoperidinoid dinoflagellates are known to be sensitive to oxidation (*80*).

#### Planktic foraminifera

Shifts in planktic foraminifera abundances have often been associated with lithological alternations, which in turn have been correlated with precession (*81*). Cold/eutrophic planktic species dominate the Late Miocene homogeneous marls deposited during insolation minima, whereas the assemblages are dominated by warm/oligotrophic species at times of insolation maxima accompanied by the deposition of laminated marls (*13*, *82*, *83*). Selli (*84*) interpreted some “dystopic faunal elements” as signs of the beginning of the salinity crisis. The most important bioevent at the Tortonian/Messinian boundary (7.25 Ma) is the replacement of right-coiled *Globorotalia menardii* form 5 by left-coiled representatives of the *Globorotalia miotumida* group (*Globorotalia miotumida* and *Globorotalia conomiozea* along with their intermediates *Globorotalia miocenica mediterranea, Globorotalia saphoae,* and *Globorotalia dalii*) (*85*). The Messinian stratigraphic distributions of the species *Globoturborotalita nepenthes*, *Globigerinoides trilobus* and neogloboquadriniids show distinct intermittent presence patterns across the basin (*13*, *83*). Unkeeled deep-dwelling globorotaliids were sparsely present after 6.72 Ma, with their disappearance being attributed to the increase of salinity at or beyond their tolerance limits (*13*). According to Kontakiotis et al.(*39*), high-salinity conditions (SSS > 40‰) were developed since 6.9 Ma in the eMed, resulting in thermal and salinity stressful conditions, which were reflected in a less diversified planktic fauna (often containing only 3–4 species) and increased abundances of taxa considered tolerant to increased salinity. This decrease in planktic foraminifera diversity started at 6.9 Ma and peaked after 6.7 Ma (*83*).

The intermittent occurrence of almost monospecific *Turborotalita quinqueloba/ Turborotalita multiloba* assemblages (*82*, *86*), and/or *Orbulina universa* blooms (*83*, *87*), and the overturn in the most abundant shallow-dwelling species (decrease of *Globigerinoides obliquus* group and increase of *Globigerina bulloides* group have been interpreted as indicative of the progressive restriction of the Mediterranean that produced a more eutrophic environment related to the salinity increase. It is worth noting that all these species can tolerate hypersaline surface waters (*88*), and particularly the small-sized opportunistic species *T. quinqueloba* and *T. multiloba* can dominate over others in such highly stressed environments (*83*). The similarly discontinuous distribution pattern of neogloboquadriniids was probably controlled by changes in the deep chlorophyll maximum, which was absent in dry periods, but expanded (though not productive) during wet periods (*13*).

#### Benthic foraminifera

Benthic foraminifera became oligospecific in the wMed already after 7.17 Ma, presumably due to changes in bottom-water oxygenation (*24*): high-oxygen benthic foraminifera species (e.g., *Siphonina reticulata*, *Cibicidoides italicus*, *C. kullenbergi*) were replaced by the species *Oridorsalis umbonatus*, *Sphaeroidina bulloides*, *Uvigerina peregrina, Gyroidinoides* spp*., Melonis* spp. (*9*, *10*, *89*), which are associated with reduced oxygen conditions and/or increased organic matter content (*90*). The benthic foraminifera records from the Po Plain- Northern Adriatic (PoA) sections Monte del Casino and Trave show analogous changes in the assemblages after 7.17 Ma (*91*). In the eMed, such changes were more intense and, in addition to the disappearance of the high-oxygen taxa, the new assemblages were dominated by buliminids (e.g., *B. elongata*, *B. subulata*, *B. aculeata*), bolivinids (*B. scalprata miocenica*, *B. dilatata*, *B. plicatella, B. spathulata*), and uvigerinids (e.g., *U. peregrina*, *U. cylindrica*, *U. striatissima*), reflecting high organic-carbon flux to the sea floor, increased salinity and/or anoxic/dysoxic conditions (*42, 92*, *93*). Benthic foraminifera communities were additionally impacted by the further deterioration of bottom-water conditions at 6.7–6.8 Ma, when deep-water ventilation was further reduced and organic carbon flux to the sea floor increased in both wMed (*24*, *83*, *94*) and eMed (*96*). Additionally, in some basins, the concomitant appearance of shallow-water species such as *Elphidium* spp., *Cibicides lobatolus*, *Discorbis* spp., *Asterigerina planorbis, Rosalina* spp., and *Valvulineria bradyana* is observed (*42*, *90*, *92*).

With the Zanclean reflooding, normal and well-oxygenated conditions were gradually reestablished in the Mediterranean basin (*96*) allowing the return of the high-oxygen species that disappeared since 7.17 Ma (e.g., *Siphonina reticulata*, *Cibicidoides italicus*, *C. kullenbergi*). In particular, the recolonization by *S. reticulata* in the Mediterranean appears to be a nearly synchronous event (*97*, *100*), recorded at the beginning of the MPI 2 biozone (*100*, *101*). The species turnover at the base of the Pliocene was stronger than the one recorded after the Tortonian/Messinian boundary (Fig. 4D).

#### Corals

Before the MSC, the most abundant genera were *Tarbellastrea*, *Solenestrea* and *Siderastrea*, while *Porites* was the main frame-builder species of pre-evaporitic Messinian reefs (*102*, *103*). Except *Solenastrea*, these genera also had the widest geographic and stratigraphic distribution suggesting that they were able to adapt to diverse paleoenvironmental conditions.

#### Bivalves

Looking only at Ostreida and Pectinida (large-sized, calcitic shells, with generally better preservation and greater stratigraphic and geographic distributions), their species richness drops from the Tortonian to the pre-evaporitic Messinian.

#### Gastropods

There is a turnover in families of medium-sized gastropods (Neogastropoda) from communities rich in species of the family Clavatulidae. In contrast, other apparent faunal changes among carnivore gastropods such as the sharp increase in number of species of the families Mangeliidae and Raphitomidae (Neogastropoda), Eulimidae (Caenogastrpopoda) and the parasitic family Pyramidellidae (Heterobranchia) in the Zanclean may be attributed to the biases of the record (see *Limitations* section in the main text) — few occurrences of these taxa in the Late Miocene, probably due to preservation and sampling bias. Moreover, the biodiversity results are driven by changes in the PoA region, because the fossil record (*26*) does not contain sufficient occurrences from all stages in all three regions to conduct the analysis.

#### Bryozoans

The Mediterranean fossil record for the Zanclean is limited to a few localities (*26*). The most abundant and diverse order of bryozoans, the Cheilostomes, is the best studied and represented in the Neogene Mediterranean deposits, corresponding to about 80% of the analyzed faunas. A relatively high proportion of cheilostome species (68%) are common between the Messinian and the Pliocene (*104*). Compared to the present-day, about 26% of present-day Mediterranean appeared before or during the Messinian, whereas 17% of the modern bryozoans in the basin are Mediterranean endemics (*104*). Apart from this marked continuity of the taxa, a progressive disappearance of thermophilic Tethyan relics has been observed (e.g., *Biflustra savarti*, *Emballotheca*, *Metrarabdotos*, *Nellia*, *Steginoporella*) and attributed to the Late Miocene global cooling, while some modern taxa of Atlantic origin only appeared in the pre-evaporitic Messinian (e.g., *Cryptosula pallasiana, Schizotheca fissa, Scrupocellaria scrupea*), mostly in the wMed (*105*) and less so in the eMed (*106*).

#### Echinoids

In the Tortonian, the Mediterranean echinoid assemblage was rich and dominated by shallow-water tropical-subtropical species, such as the genera *Clypeaster* and *Echinolampas* (*107*).

The stressful environmental conditions in the pre-evaporitic Messinian would be expected to have an impact on echinoids, which are exclusively marine invertebrates, intolerant of great salinity fluctuations that are never found in freshwater (*108*) and rarely can tolerate moderate salinity changes (*109*).

#### Bony fishes

Otolith and skeletal findings were combined to obtain the Tortonian–Zanclean fossil record of bony fishes (*26*), which collectively represent both the shallow and deep environments before and after the MSC within the Mediterranean and the three regions. The Tortonian and pre- evaporitic Messinian bony fish record of the Mediterranean (*26*) comprised species of Mediterranean–Atlantic distribution (*110*) and several with Paratethyan-affinity (*111*, *112*). In the pre-evaporitic Messinian, the fossil record shows the extirpation of several pelagic species that were common in the Mediterranean in the Tortonian, including mesopelagic fishes of the family Myctophidae such as *Benthosema fitchi*, *B. glaciale, Lobianchia gemellarii, Myctophum punctatum, Notoscopelus bolini* and *N. resplendens* and deep-water fishes such as *Coelorinchus* spp., *Trachyrincus* spp. and *Verilus* spp. (*111*, *113*, *114*). Notably, after 6.8 Ma, otolith isotopic data have indicated that benthic fish growth was severely hampered by a combination of high salinity stratified bottom waters with high temperature fluctuations, leading to their final disappearance from the sea bottoms at least in the eMed, even before the MSC onset (*94*). All of these species re-established in the Mediterranean after the MSC. Additionally, several endemic Mediterranean species appeared during the pre-evaporitic Messinian, e.g. myctophids such as *Ceratoscopelus dorsalis* and *C. miocenicus* (*115*, *116*) and benthic gobies such as *Buenia affinis* and *Caspiosoma lini* (*112*, *115*); most, but not all, of these endemics disappeared in the Zanclean. Apart from the reintroduction to the Mediterranean of common Tortonian species in the Zanclean, we also observe the first appearance in the basin of common extant benthic species *Chromogobius zebratus, Gobius guerini, Gobius geniporus, Gobius niger, Lesueurigobius sanzi, Zebrus zebrus* and *Zosterisessor ophiocephalus* (*112*, *117*, *118*).

#### Sharks

The scarcity of Messinian records of elasmobranchs complicates the reconstruction of the Late Miocene Mediterranean fauna (*26*). That said, the Messinian assemblage from Algeria (*119*) strongly resembles other central Mediterranean Tortonian assemblages (e.g., *120*, *121*) in terms of taxonomic composition. Most of the Zanclean assemblage is represented by relic Miocene taxa. In the lowermost Pliocene, published occurrences taxa like *Carcharocles megalodon* and *Hemipristis serra* hint at the persistence of the Miocene structure of the Mediterranean fauna during the Early Zanclean (*122*). Other relic Miocene species (e.g., *Megascyliorhinus miocaenicus*, *Pachyscyllium dachiardii*, *Pachyscyllium distans*, *Cosmopolitodus plicatilis* and *Parotodus benedeni*) even persisted in the Mediterranean until the mid-Pliocene at least (*123*, *124*). In the Late Miocene and Early Pliocene, *Carcharhinus* was likely the most speciose shark genus in the Mediterranean. The diverse appearance of the Pliocene *Carcharhinus* assemblage is seemingly due to the in-depth revision of the Pliocene carcharhines provided by Marsili (*125*). On the other hand, most Miocene carcharhines were attributed to the ‘wastebasket’ fossil species *Carcharhinus priscus* and *Carcharhinus egertoni* (*126*). While the former has been recently redefined on the basis of Agassiz’ types by Reinecke et al. (*127*), a comprehensive revision of the Miocene Mediterranean teeth of *Carcharhinus* is still needed.

No strong breaks other than those observed globally can be recognized for the Mediterranean elasmobranch faunas around the Miocene/Pliocene transition. The lowermost Pliocene nearshore assemblages (*122*, *128*) are, at various degrees, reminiscent of the Miocene faunas.

#### Marine mammals

The Late Miocene–Early Pliocene fossil record of marine mammals of the Mediterranean includes representatives of cetaceans (whales and dolphins), pinnipeds (seals) and sirenians (dugongs). The cetacean Tortonian Mediterranean record is quite diverse both among the echolocating toothed whales (Odontoceti) and the baleen whales (Mysticeti). Odontocetes are represented by Delphinida (dolphins), Physeteroidea (sperm whales) and Ziphiidae (beaked whales); while at least four mysticete families are present: Balaenidae (right whales), Balaenopteridae (rorquals), Eschrichtiidae (gray whales) and Cethotheriidae. The fossil record of cetaceans from the pre-evaporitic Messinian of the Mediterranean is limited to three specimens belonging to odontocetes, including a Phocoenidae (porpoise) and a Physeteroidea. The Zanclean cetacean fauna consists of almost the same families as the Tortonian fauna, but the Early Pliocene brings greater diversity at the genus and species levels, which could, however, be attributed to the better preservation of the Pliocene specimens.

All determinable fossil remains of sirenians in the Late Miocene–Early Pliocene Mediterranean belonged to dugongid genus *Metaxytherium* represented by three putative chronospecies: the medium-sized *M. medium* (Tortonian), the small-sized *M. serresii* (latest Tortonian–earliest Zanclean), and the large-sized *M. subapenninum* (Zanclean). Although the fossil record of pinnipeds from Miocene Mediterranean deposits is extremely fragmentary (*129*), the continued presence of the seals Monachinae in the Mediterranean from the Early Miocene to the present day can be attested, maintaining low diversity and small, geographically distinct populations. Based on the Messinian record, only represented by *Messiphoca mauritanica* from Algeria, it can be speculated that, before the MSC onset, pinnipeds were confined to even more restricted areas, most or all of which close to Gibraltar, a scenario similar to the one observed today for *Monachus monachus* (*130*).

---

**Fig. S1.**
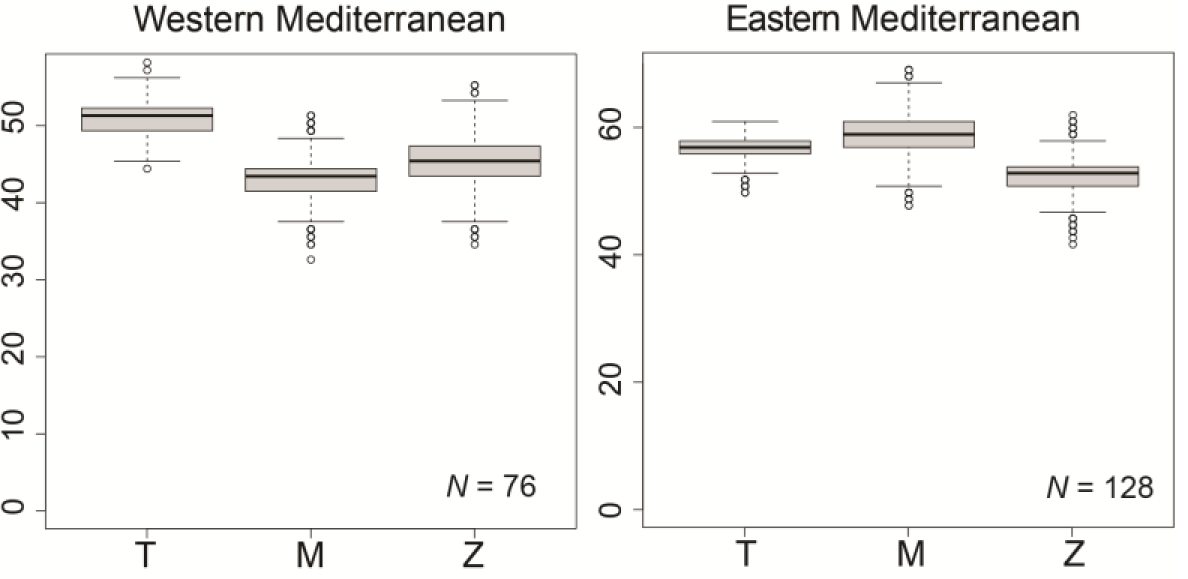
Calcareous nannoplankton species richness through time in the Western and Eastern Mediterranean across the Tortonian (T), pre-evaporitic Messinian (M) and Zanclean (Z).

**Fig. S2.**
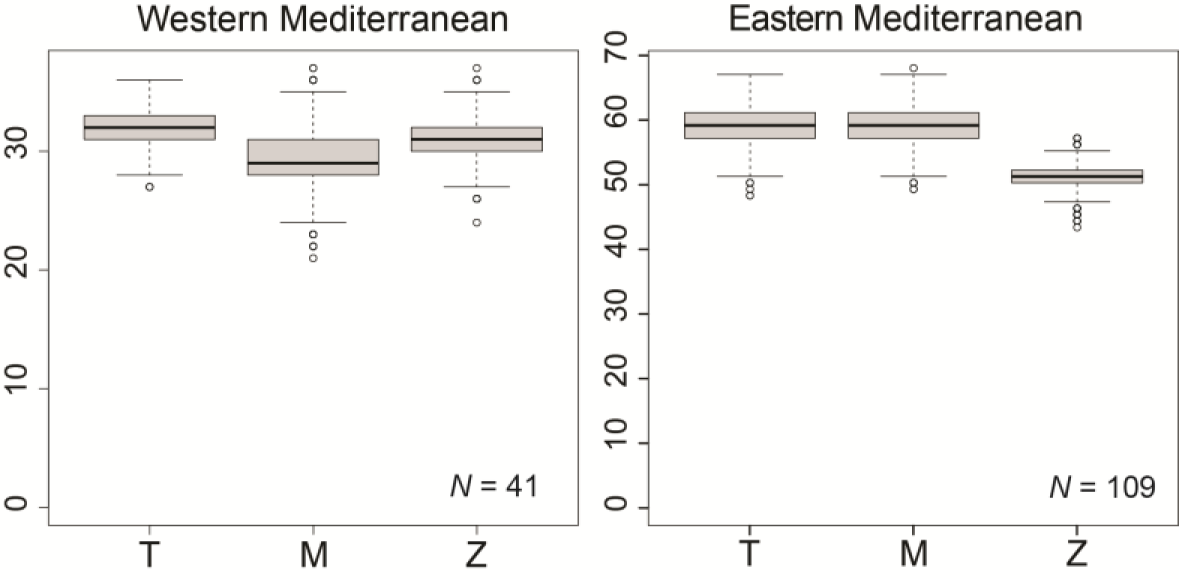
Dinocyst species richness in the Western and Eastern Mediterranean across the Tortonian (T), pre- evaporitic Messinian (M) and Zanclean (Z).

**Fig. S3.**
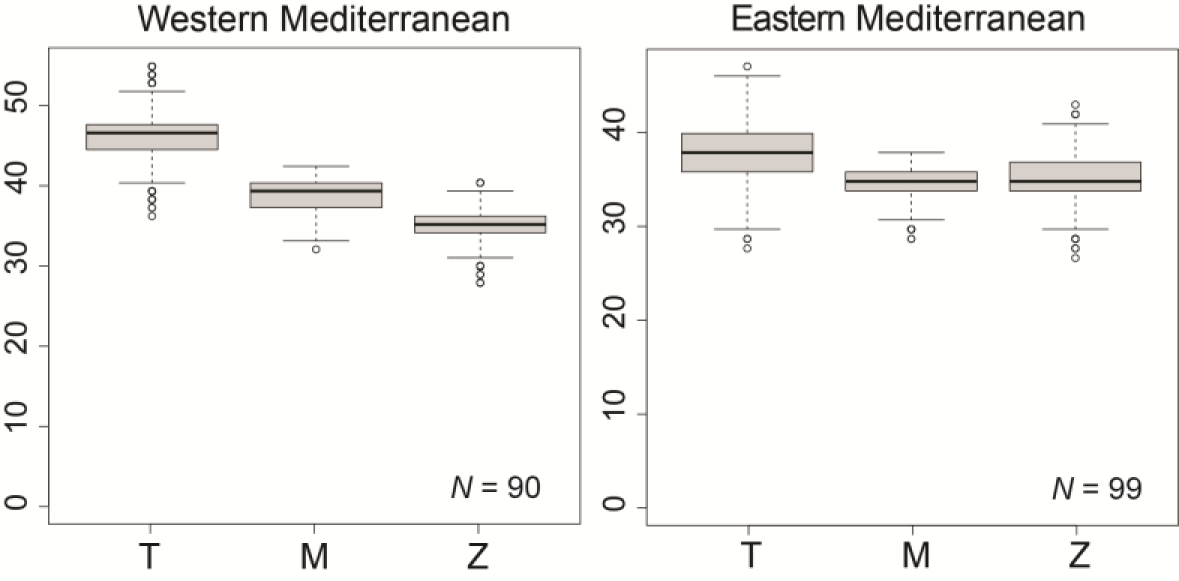
Temporal and spatial changes in species richness of the planktic foraminifera in the Mediterranean regions in the Tortonian (T), the pre-evaporitic Messinian (M), and the Zanclean (Z).

**Fig. S4.**
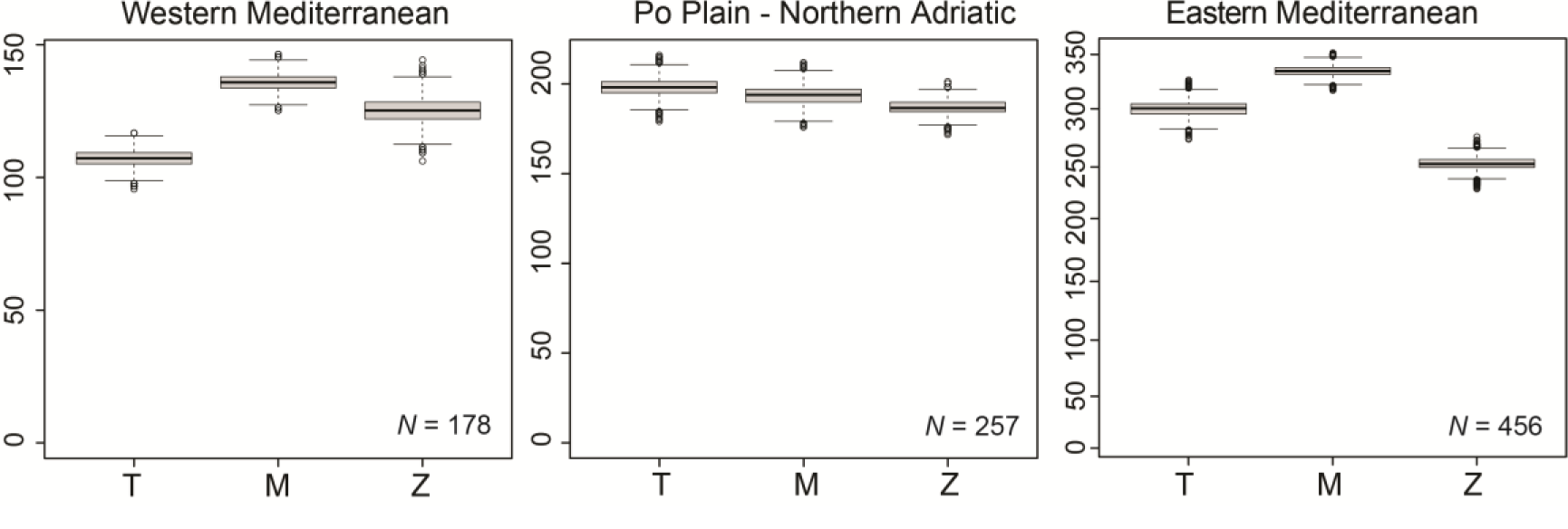
Ostracod species richness in the Mediterranean regions in the Tortonian (T), the pre-evaporitic Messinian (M), and the Zanclean (Z).

**Fig. S5.**
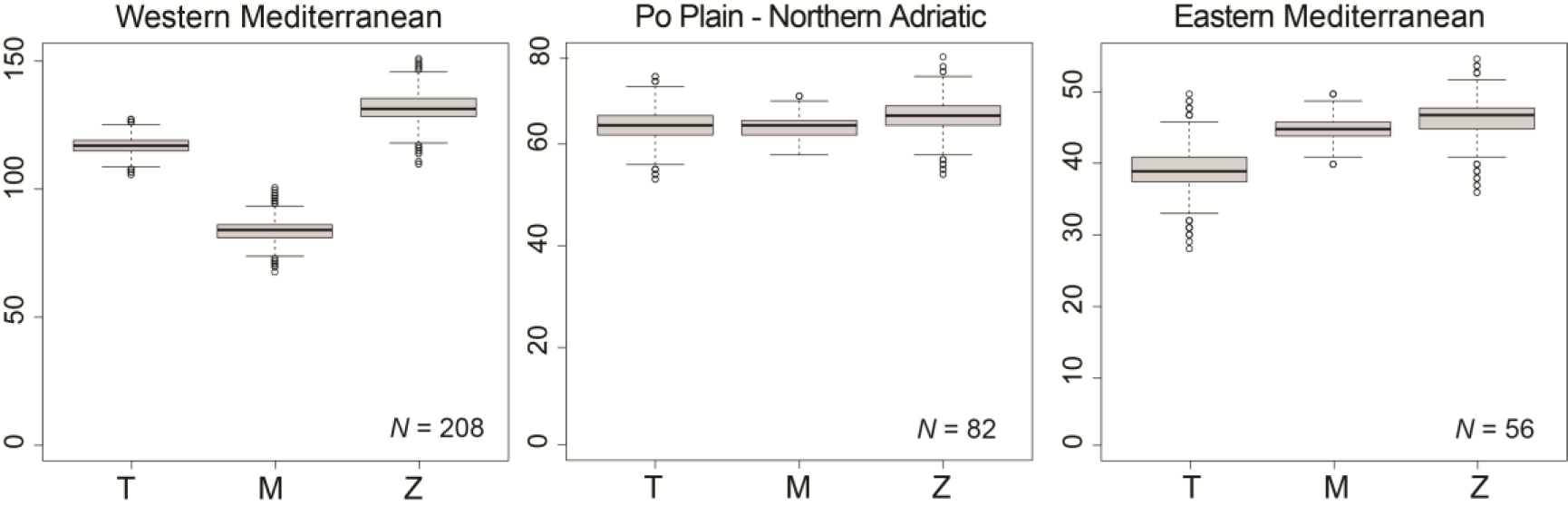
Changes in bivalve species richness of the Mediterranean Sea from the Tortonian (T), the pre-evaporitic Messinian (M) to the Zanclean (Z).

**Fig. S6.**
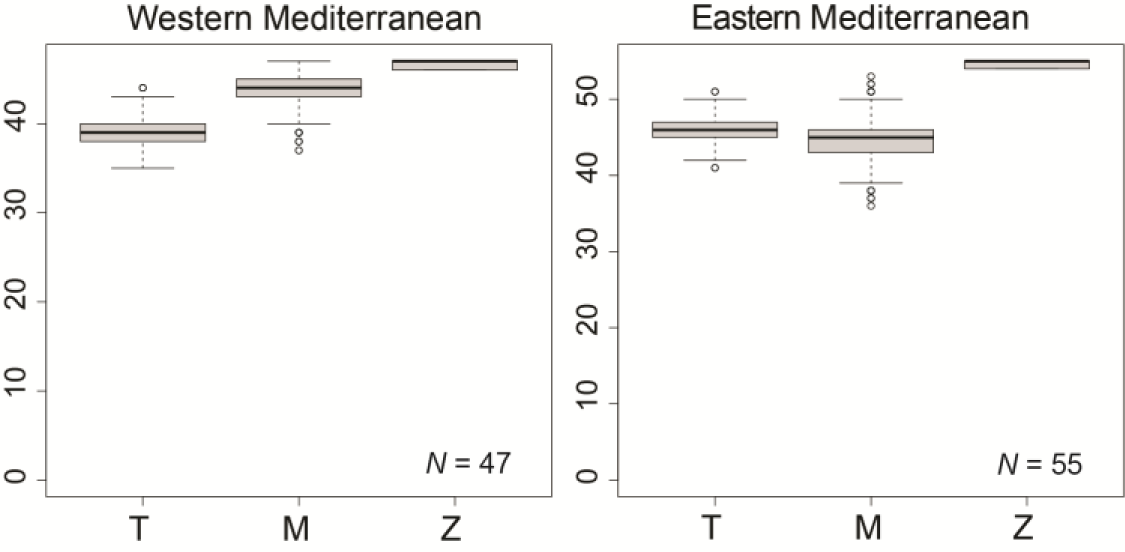
Species richness of the bryozoan fauna in the Western and Eastern Mediterranean in the Tortonian (T), the pre-evaporitic Messinian (M) and the Zanclean (Z).

**Fig. S7.**
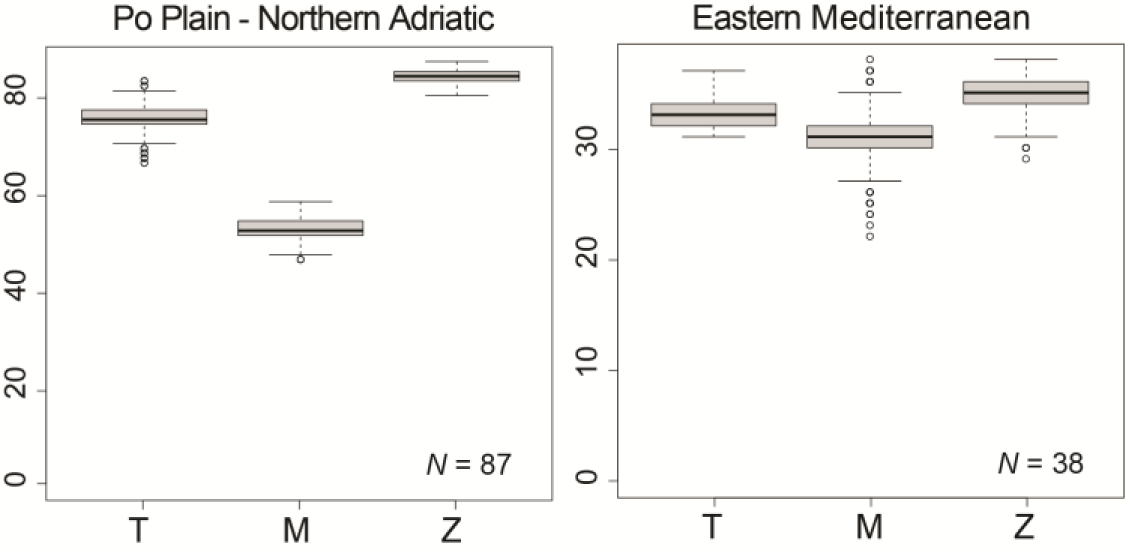
Species richness of bony fishes within the Eastern and the Po Plain-Northern Adriatic region in the Tortonian (T), the pre-evaporitic Messinian (M), and the Zanclean (Z).

**Table S1.** Summary of results obtained from bibliographic data for organic-walled dinoflagellate cysts for the Mediterranean and the three time intervals considered.

|  | West |  |  | Po Plain-Adriatic |  |  | East |  |  |
| --- | --- | --- | --- | --- | --- | --- | --- | --- | --- |
|  | T | M | Z | T | M | Z | T | M | Z |
| Sequences | 3 | 5 | 3 | 2 | 2 | 1 | 10 | 13 | 13 |
| Species | 53 | 77 | 52 | 41 | 42 | 21 | 90 | 92 | 73 |
| Genera | 24 | 25 | 20 | 24 | 23 | 11 | 38 | 37 | 26 |
| Gonyaulacoid species | 39 | 50 | 27 | 29 | 29 | 14 | 57 | 56 | 53 |
| Protoperidinioid species | 4 | 1 | 6 | 10 | 8 | 1 | 10 | 17 | 8 |
| Goniodomacean species | 8 | 8 | 2 | 3 | 3 | 2 | 4 | 4 | 3 |
| Protoperidin/Gonyaulacoid ratio | 10% | 2% | 18% | 26% | 22% | 7% | 15% | 23% | 13% |

